# Aurora B inhibition promotes a hyper-polyploid state and continued endomitotic cycles in RB and p53 defective cells

**DOI:** 10.1101/2024.03.27.585450

**Authors:** Shivam Vora, Ariel Andrew, Ramyashree Prasanna Kumar, Deborah Nazareth, Madushan Fernando, Mathew JK Jones, Yaowu He, John D. Hooper, Nigel AJ McMillan, Jelena Urosevic, Jamal Saeh, Jon Travers, Brian Gabrielli

## Abstract

Polyploidy is a common outcome of chemotherapies, but there is conflicting evidence as to whether this is a source of increased chemotherapy resistance and aggressive disease, or a benign or even favorable outcome. We have used Aurora B kinase (AURKB) inhibitors that efficiently promote polyploidy in many cell types to investigate the fate of polyploid cells. We demonstrate AURKB inhibitor treatment of cells that have loss of RB and p53 function causes them to become hyper-polyploid, undergoing continuous rounds of growth, replication and failed mitosis/cytokinesis (endomitosis), whereas RB and p53 functional cells will eventually exit the cell cycle. These hyper-polyploid cells (>4n DNA content) are viable and undergo continuous endomitotic cycles, but have lost the ability to form viable colonies <u>in vitro</u> or form tumours <u>in vivo</u>. Investigation of mitosis in these cells revealed that centrosome duplication remained coupled to DNA replication, with the hyper-polyploid cells containing high numbers of centrosome that were capable of supporting functional mitotic spindle poles, but these failed to progress to anaphase/telophase structures even when AURKB inhibitor was removed after 2-3 days. However, when AURKB inhibitor was removed after 1 day and cells had failed a single cytokinesis to become tetraploid, they retained long term colony forming ability. Collectively, these findings demonstrate that tetraploidy is well tolerated by tumour cells but higher ploidy states are incompatible with long term proliferative potential.

## Introduction

Polyploidy defines any cell with >2n DNA content, the most common forms of polyploidy observed is tetraploid (4n) or aneuploid usually 2-4n DNA content. Mechanisms that promote polyploidy are endoreplication, where cells undergo rounds of replication without an intervening mitosis, endomitosis, where cells undergo a normal cell cycle but fail cytokinesis resulting in either a single enlarged nucleus, binuclear or multinuclear phenotypes depending on which components of cytokinesis are defective, or cell fusion [1].

Polyploidy is commonly characterized by whole genome duplication events which lead to an increase in size and metabolic capacity of the cell or organism. Polyploidy is a consequence of aberrant cell division which may result from environmental stressors or in some cases by physiological drivers. The prevalence of polyploidization differs greatly based on the class and kingdom. In plants where polyploidy was first discovered, it is extremely common whereas in animals it is quite rare. In humans, programmed polyploidy is observed in several instances within organs such as the liver (hepatocytes), bone marrow (megakaryocytes), placenta (trophoblast giant cell), skeletal muscles and osteoclasts, and also in tissue regeneration. In the context of human cancers, polyploid cells are present in cancer as well as in premalignant lesions but their role is less well understood than in physiologically programmed polyploidy [2]. Conflicting studies have demonstrated the tumorigenic potential of polyploid cells, although this is thought be through a process of depolyploidisation where a daughter cell is produced with reduced ploidy that has inherited increased fitness{Mosieniak, 2015 #94} [3].

The best studied models for polyploid somatic cells in tissue regeneration are hepatocyte development and liver regeneration. Polyploidisation is a common feature of liver development and in response to liver damage. This polyploidy normally featured tetraploid and octaploid cells, generally the product of failed cytokinesis. These polyploid hepatocytes can proliferate and also undergo reductive divisions to produce diploid daughters [4]. The physiological role of tetraploid cells is well demonstrated in a mouse model of hepatocyte-specific deletion of CDK1. Partial hepatectomy results in massive hepatocyte proliferation, but in the CDK1-deleted model cells can undergo a single round of replication but not division, resulting in massive tetraploidy. Normal liver regeneration was observed in the CDK1-deleted model indicating that the tetraploid cells were capable of regeneration of liver bulk and function [5–7]. This also occurs normally in liver, but the normal hepatocytes can then revert to 2n by reductive division [4]. Other tissues use similar programs for repair and maintenance of homeostasis [8].

Genome duplication is a common feature of cancers [9]. This has been proposed to provide buffer for changes in gene expression, mutations and deletion [10]. There is an extensive literature on polyploid giant cancer cells (PGCC), a commonly observed feature cultures of cancer cell lines and observed *in vivo*. As with physiological polyploidy, the PGCCs are the result of either cell fusions, endoreplication or failed cytokinesis, and are a common result of chemotherapy treatments. The consensus view is that PGCC are a source of more aggressive and treatment resistant tumours [1, 8, 11–14].

Several studies have investigated the formation and significance of polyploid giant cancer cells (PGCC) in response to chemotherapy drugs. The contribution of these cells in cancer is unclear due to conflicting evidence of their tumour suppressive or tumorigenic potential. Studies have reported that PGCCs are more likely to undergo apoptosis [15] and enter therapy induced senescence [16], whereas several studies have shown the ability of tetraploid cells to give rise to normal progeny which may promote cell survival through an enhanced resistance to therapy [17] {Puig, 2008 #100}. Senescence can be considered a favourable therapeutic outcome as it is an irreversible cell cycle arrest that safeguards against tumour progression {van Deursen, 2014 #105}. Inactivation of tumour suppressors p53 and RB that are essential in the senescence pathway is a common feature of cancers bypasses senescence [18].

Here we have investigated the polyploidy driven by AURKB inhibition. AURKB inhibitors (AURKBi) have been investigated in a broad range of cancers, but as yet none have been approved for clinical use [19]. Many of the developed AURKi have activity towards all three Aurora kinases, AURKA, AURKB and AURKC, although AURKA and AURKB are the primary targets in cancers [20]. AURKA and AURKB are functionally distinct regulators of progression through mitosis. AURKA is essential for centrosome maturation and regulates mitotic entry. AURKB regulates exit from mitosis and controls correct partitioning of the replicated genome [21]. Inhibition of AURKB disrupts normal chromosome alignment and segregation during mitosis, premature inactivation of the spindle assembly checkpoint-dependent mitotic arrest and cytokinesis failure resulting in polyploidy [22–24]. This polyploidy triggers a p53-dependent cell cycle arrest through HIPPO pathway activation to block endoreplication [25].

AURKB selective inhibitors promote a failed cytokinesis, whereas the dual inhibitors cause a modest mitotic delay then failed cytokinesis [23, 24, 26]. The outcomes of AURKBi treatment appear to be cell line dependent; cell proliferation may be arrested, or cells may become senescent or undergo apoptosis [22, 26–31]. Senescence is the primary response to AURKB inhibition in cells containing wild type RB and p53, but cells with defective RB and p53 such as mutation/deletion or inactivation by viral oncogenes such as HPV E6/E7 bypass this senescence arrest (Vora et al., submitted). Loss of RB has also been reported to increase sensitivity to AURKB inhibitors [24, 29, 32]. Here we have investigated the outcomes of AURKBi treatment in RB and p53 defective tumour cell lines. We demonstrate they efficiently promote the formation of polyploid giant cells, and demonstrate that although they retain viability they have lost long term proliferative potential compared with tetraploid cells that retain long term proliferative potential.

## Results

### Loss of RB1 function does not effect acute sensitivity to AURKBi in tumour cell lines

Several recent studies have reported that RB defects are critical for sensitvity to AURK inhibitors [26, 32]. To determine how universal this effect was, we have analysed the Cancer Dependency Map (DepMap [33]) dataset for sensitivity to AURKA/Bi Alisertib and the AURKBi AZD2811 (Barasertib in the data set) from the Cancer Target Discovery and Development (CTD^2 [34]) and Genomics of Drug Sensitivity (GDSC1/2 [35]) datasets that together have sensivtiy data for over 1000 human cancer cell lines. 11% of CTD^2 cell lines contained *RB1* mutants, the majority being destablising mutant resulting in low RB protein levels (Supp. Figure S1A). There was no significant difference in the range of sensitivities of the *RB1* mutant and wild type containing cell lines to either AZD2811 or Alisertib (Supp. Figure S1B). The lack of effect was not due to significant difference in the doubling time of the RB wild type and mutant cell lines that would be expected to effect the activity of mitotically-targeted agents such as AURKi (Supp. Figure S1C). Similar was observed with Alisertib in the GSCD1 dataset. These data suggest that loss of function of RB was not responsible for sensitivity to AURKBi, at least in terms of acute effects on viability, but it may influence the long term viability of RB mutant cells. RB mutations are rare in the absence of p53 mutation (TCGA Pan Cancer, CCLE; Supp. Figure 1D), therefore the effects of longer term AURKBi treatment on RB+p53 wild type and RB+p53 defective (defective for both RB and p53) tumours was investigated.

### AURKBi promote hyper-polyploidy in RB+p53 defective cells

The role of RB and p53 in limiting polyploidy triggered by AURKB inhibition was investigated in a panel of cancer cell lines, either with functional RB and p53 (HT1080) or loss RB and p53 funciton (RB+p53 defective) through mutation (C33A) or due to HPV E6/E7 expression (CaSki). Treatment with the AURKBi AZD2811 for 3 days had no apparent effect on the replicative abiltiy of the RB and p53 defective C33A and CaSki cells lines demonstrated byEdU incorporation, whereas RB and p53 wild type HT1080cell line had significantly reduced replicative activity (Figure 1A). The replicating C33A and CaSki cells were hyper-polyploid (>8n DNA content; Figure 1B, Supp. Figure S2A). Even with 6 days AZD2811treatment there was little reduction in either EdU incorporation or immunostaining for Ki67, another marker of proliferation (Figure 1C). The cells continued to increase in size, the increased nuclear size correlated with the increased cell size (Supp. Figure 2B). Even after 2 days treatment, the nuclei were multilobed often with the presence of micronuclei and by 6 days treatment, the nuclei were commonly fractured (Figure 1D). The multilobed phenotype observed with 2 days treatment was also observed in RB and p53 wild type HCT116 cells, but unlike the RB and p53 defective cells, the size and phenotype were unchanged with 6 days treatment (Supp. Figure S2C,D). Ki67 staining was dramatically reduced with 2 days treatment in the wild type cells, indicating that AZD2811 treatment induced cell cycle arrest with 2 days treatment, with almost complete loss of Ki67 staining by 6 days. Engineered HCT116 RB^−/−^ p53^−/−^ cells treated with AZD2811 for 6 days also retained Ki67 staining and continued to increase in size (Supp. Figure S2D,E). This demonstrated that loss of RB and p53 function was critical to bypassing AURKBi-induced cell cycle arrest.

**Figure 1.**
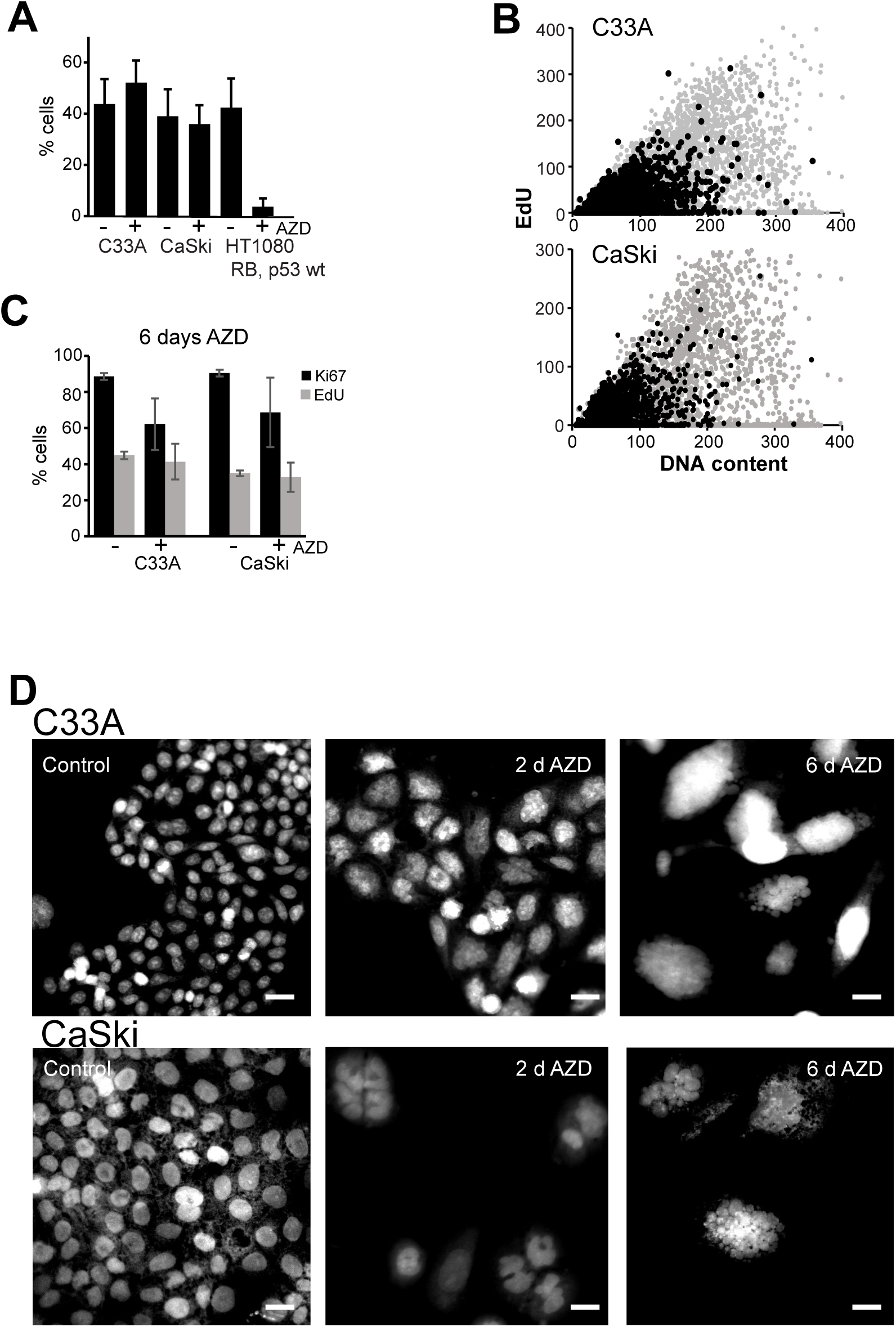
**A**: RB/p53 wild type HT1080 and RB/p53 defective C33A and CaSki cells were treated for 3 days with and without (control) 200 nM AZD2811 (AZD; AURKBi) then labelled for 2 h with EdU and stained for EdU incorporation. The percentage EdU positive cells was assessed in triplicated samples. **B**: C33A and CaSki cells were treated with 200 nM AZD2811 for 2 days then EdU labelled for 2 h, fixed and stained for EdU and DNA, then analysed by high content imaging. Control cells are shown as the black dots and AZD2811 treated as the grey dots. >4000 cells were analysed for each condition. **C:** C33A and CaSki cells were treated with 200 nM AZD2811 for 6 days then EdU labelled for 2 h, fixed and stained for EdU and Ki67, then analysed by high content imaging. The data are the mean and standard deviation of triplicate wells. **D** C33A and CaSki cells were treated with 200 nM AZD2811 for 2 and 6 days, fixed and stained for DNA, then imaged. The bar indicated 20 μm.

The efficiency of the AZD2811 in bypassing cytokinesis was assessed by time lapse imaging of C33A and CaSki cells treated with AZD2811 over 3 days. Over this time, control cells undergo 3-4 mitoses, >90% producing two daughter cells in cytokinesis (Figure 2A,B). The AZD2811-treated cells all failed cytokinesis up to 4 times across the 3 days (Figure 2B,C). The effects of AZD2811 were due to inhibition of AURKB as two other AURK inhibitors, alisertib and AMG900 that inhibit both AURKB and AURKA, produced similar failure of cytokinesis over the same 3 day period (Figure 2D).

**Figure 2.**
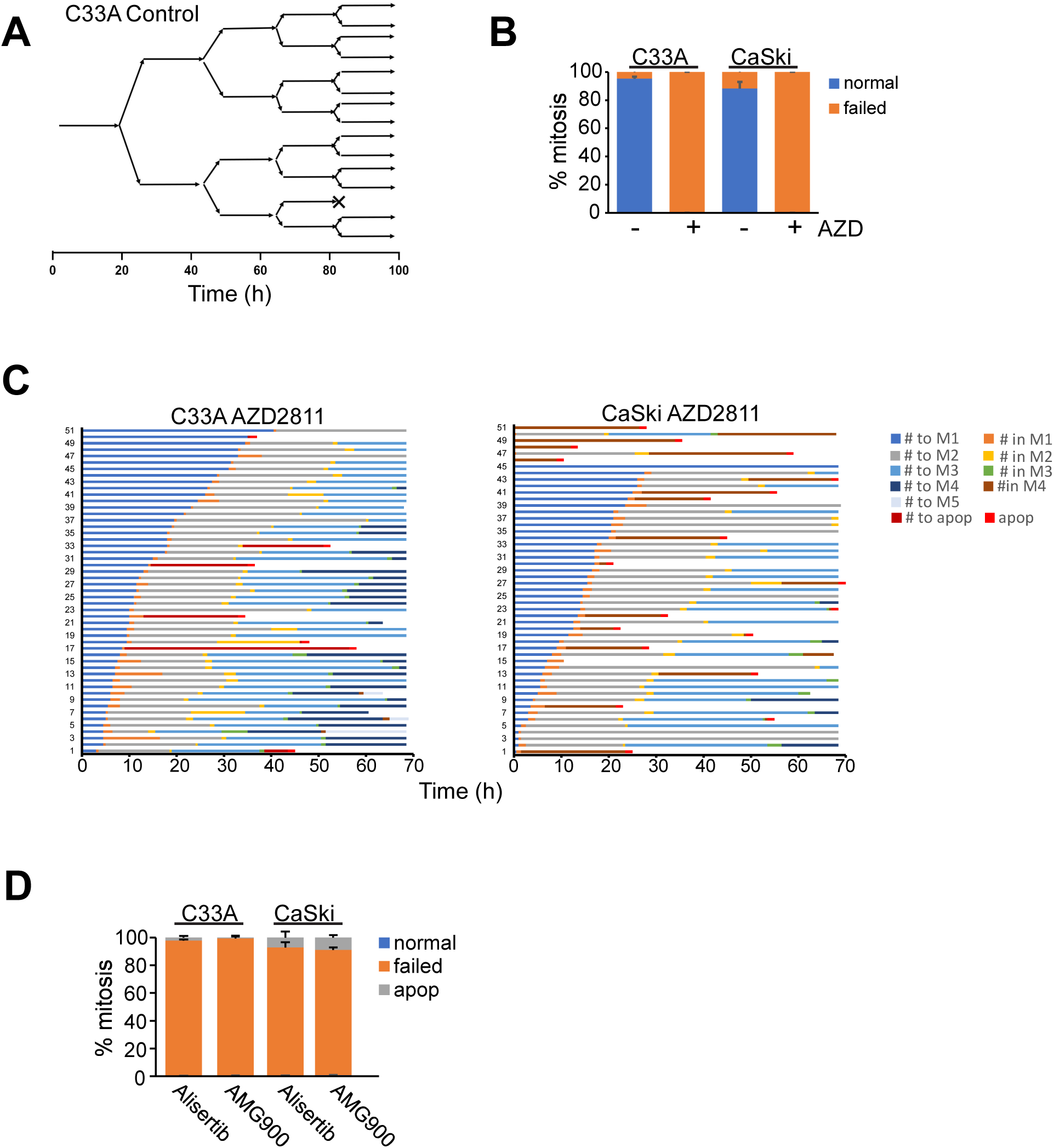
**A**: Timeline of a single C33A cell followed for 3 days using time lapse microscopy. It shows that the majority of mitosis and cytokinesis resulted in two daughter cells. **B:** The percentage of normal and failed cytokinesis from time lapse experiments of C33A and CaSki cells without and with 200 nM AZD2811 treatment followed for 3 days. The data are from >50 cells for each condition. **C:** Timelines for C33A and CaSki cells treated with AZD2811 as in **B** and followed for 3 days. **D:** C33A and CaSki cells treated with either 1 μM alisertib or 300 nM AMG900 and followed for 3 days by time lapse microscopy. The percentage of normal and failed cytokinesis was determined for >50 cells.

The failed cytokinesis appeared not to result in excessive DNA damage. Staining for γH2AX revealed a modest increase with brightly stained foci mostly representing micronuclei associated with fractured nuclei of the hyper-polyploid cells (Supp. Figure S3). This did not appear to be restricted to only RB+p53 defective cells and was observed in all cell lines driven into polyploidy with AURKBi treatment.

### Hyper-polyploid cells are incapable of producing proliferative subclones

The high efficiency with which AURKBi promote the hyper-polyploid state raised question about whether hyper-polyploid cells had the ability to undergo reductive mitosis to produce near diploid or even tetraploid daughter cells that possessed proliferative capacity. Time lapse imaging was performed with C33A and CaSki cells that had been treated for 2-3 days with AZD2811, then drugs washed off and cells imaged from day 6-10 after commencement of treatment. Control cells proliferated normally, whereas AZD2811-treated cells continued to progress through endomitotic cycles and fail cytokinesis (Figure 3A-C; Supp. Movie 1-3). Despite the efficiency of AURKBi to inhibit cytokinesis, there was abundant evidence of large cells undergoing cytokinesis to produce 2-3 daughter cells through several rounds of mitosis (Figure 3B, arrowhead; Figure 3C yellow arrowhead). However, these apparently viable clones were a relatively uncommon feature. The ∼40% of the relatively normal cytokinesis was contributed by the small colonies of large cells undergo several rounds of cell division (e.g. Figure 3B, arrowhead). Most cells undergoing mitosis failed cytokinesis (∼40%) or had an abnormal cytokinesis (10-20%). This produced either three or more relatively equally sized daughters, or cells underwent asymmetric division producing one large and one much small daughter cells, the common outcome of this was death of both daughters (Figure 3C, white arrowhead; Figure 3D). Many of the very large multinuclear cells also died during imaging (Figure 3C, red arrowhead). Despite cell continuing to undergo mitosis, the confluence of cultures rarely increased during imaging when compared to control cells, a combination of failed cytokinesis and cell death (Supp. Movie 4). This was confirmed by colony forming assays with CaSki and C33A cells that revealed no colonies formed with only three days AURKBi treatment, although large individual cells were observed (Figure 3E). In another model, immortalised human keratinocytes expressing HPV E6/E7 similarly failed to form colonies with 3 day AURKBi treatment (Figure 3F).

**Figure 3.**
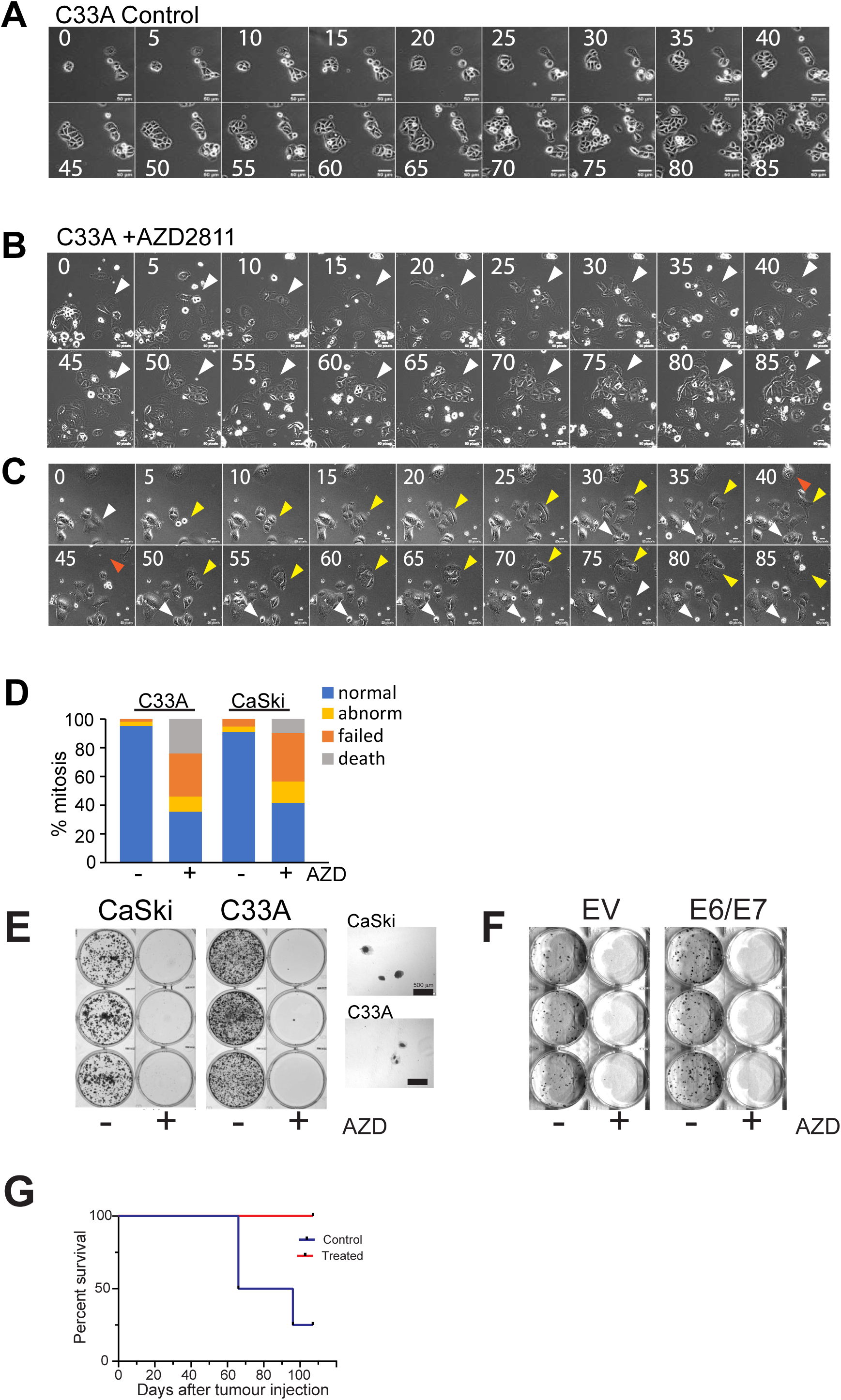
Time lapse images of **A:** control C33A cells, **B** and **C:** treated with 200 nM AZD2811. Cells were treated for 3 days then drug removed, and cells imaged from day 4 – 8. Images were collected every 30 min, and montage shows every 5 h. **D.** Outcomes of cytokinesis from the time lapse shown in A-C and a parallel experiment using CaSki. Each condition is from 100 cells. **E:** Colony forming assays of the indicated cell lines treated for 3 days with AURKBi then washed out and cells allowed to form colonies. **F:** Similar experiment to **E** using immortalised human keratinocytes (EV) or expressing HPV E6/E7. G. Nude mice were injected with either control or 5 day AZD2811 treated HCT116 RB^−/−^p53^−/−^ and mice followed. Mice were culled when tumours were >100 mm^3^ and conformed by autopsy.

To further investigate whether the AURKBi-induced giant polyploid cells can generate viable daughter cells *in vivo*, parental HCT116 RB^−/−^p53^−/−^ cells, and giant cells derived from 5 days AZD2811 treatment were injected into nude mice for an *in vivo* tumour formation assay. 3 million parental or AURKBi giant cells were injected in flanks of nude mice (the giant cells had to be injected into two sites due to their >10 fold large cell volume) and tumour growth monitored. Despite the long tumour growth delay, parental tumour growth was observed in 3 of 4 controls by 107 days, but no tumour formation was observed in the AURKBi giant cells injected mice even after 107 days (Figure 3G). Autopsy of the mice failed to reveal tumour growth at any sites within the mice.

One possible outcome of the failed cytokinesis was a disconnection between the cell replicative cycle and centrosome cycle, resulting in a relatively small number of centrosomes allowing cells to establish-relatively normal spindles despite the large numbers of chromosomes (>8n). Alternatively, centrosome duplication was not affected but the centrosomes cluster to form a smaller number of spindle poles permitting normal appearing cytokinesis [36, 37]. Either of these may explain the 40% of relatively normal appearing cytokinesis of even very large cells, and the many attempted and failed cytokinesis observed in these large cells (Supp. Movies 2-4). When cells were stained for centrosomes and spindle formation using γ-tubulin and α-tubulin antibodies, control cells displayed the expected bipolar spindle with centrosomes forming the apexes of the mitotic spindle (Figure 4A). The large cells after 6 days AZD2811 treatment were found to have multiple centrosomes usually clustered centrally in interphase cells, but these dispersed to form individual spindle poles in mitosis, although occasional clusters of two centrosomes in a single spindle pole were observed (Figure 4B,C). The centrosomes were spaced throughout the cells (Supp. Movie 5). Counting γ-tubulin stained centrosomes in mitotic spindle poles confirmed the increased centrosome numbers. The majority of control cells had two centrosomes, whereas by 2 days treatment, centrosome numbers have increased to means of between 4 and 11, depending on the proliferation rate of the cell line (CaSki was slowest and HCT116 the fastest; Figure 4D,E). Interestingly, the few mitotic cells found in the HCT116 wild type cells (RB and p53 wild type) had a mean of 8 centrosomes, suggesting these cells had undergone two rounds of failed replication and mitosis/cytokinesis. By 6 days treatment, C33A cells had an average of 32 centrosomes, whereas the few remaining attached CaSki and HCT116 RB^−/−^p53^−/−^ cells had average of 23 and 38 respectively. No mitotic cells were found in the HCT116 wild type cells at day 6.

**Figure 4.**
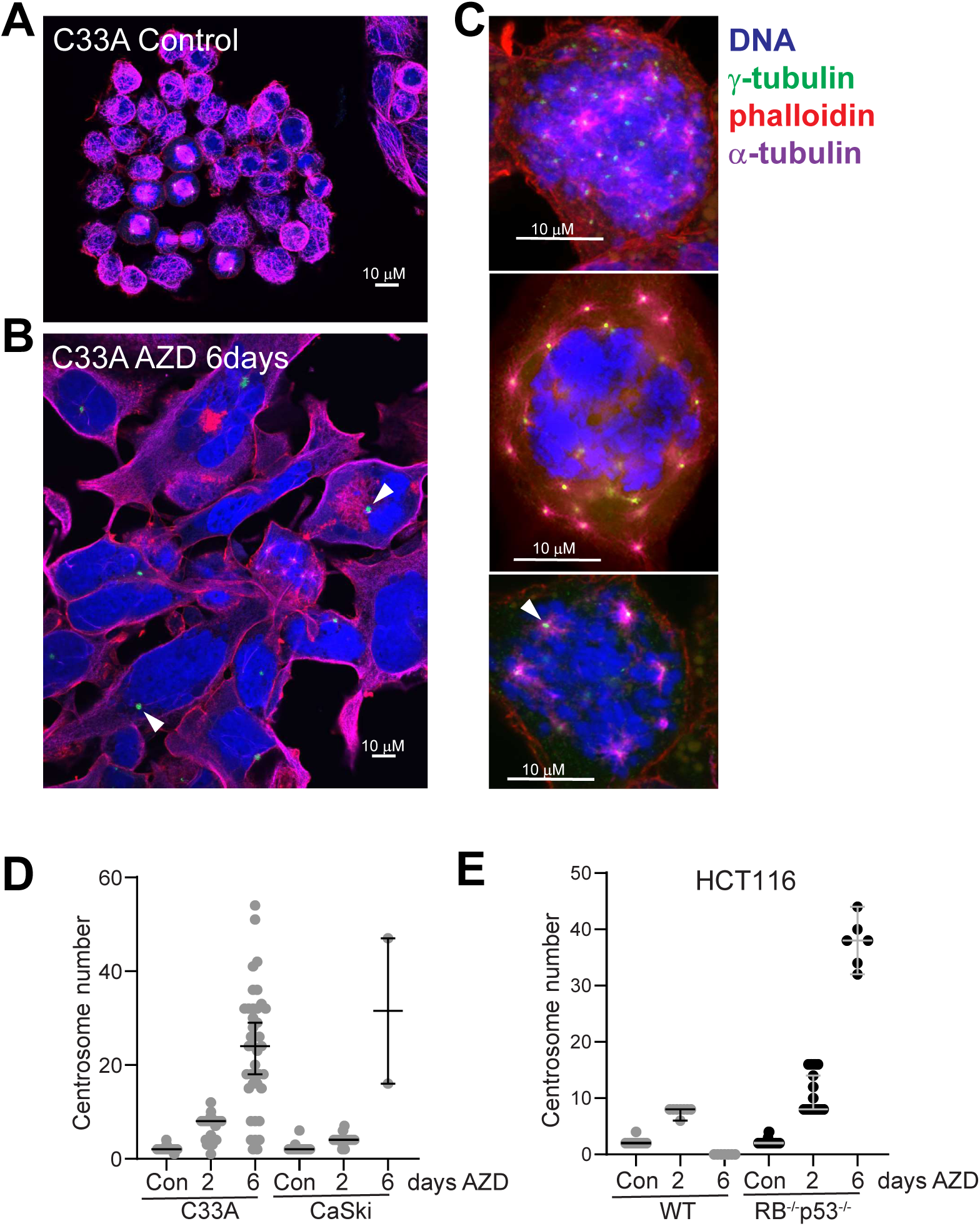
**A**: Confocal microscopy of C33A cells stained for DNA, γ-tubulin, α-tubulin to mark centrosomes and mitotic spindles and phalloidin for the actin cytoskeleton. **A:** Control, **B:** 6 day 200 nM AZD2811 treated interphase and **C:** mitotic cells. The bar in each is 10 μm. **D:** C33A and CaSki cells treated with 200 nM AZD2811 for the indicated times and stained as in **A-C** were visually inspected and centrosome number of the mitotic cells at each time point counted. **E:** Parallel experiment to **D** but using either HCT116 wild type or RB^−/−^p53^−/−^ cells.

Analysis of the 6 day treated cultures revealed that the very few colonies of smaller cells were unlikely to represent normal control cells, due to their larger size and commonly increased number of centrosome/spindle pole (Figure 5A). Some very large mitotic cells were also found to form asymmetric mitotic cells, which were likely to be forerunners of the asymmetric cytokinesis observed in the time lapse sequences (Figure 5B; Supp. Movie 3). There was also evidence of budding of small buds that contain little or no DNA (Figure 5C).

**Figure 5.**
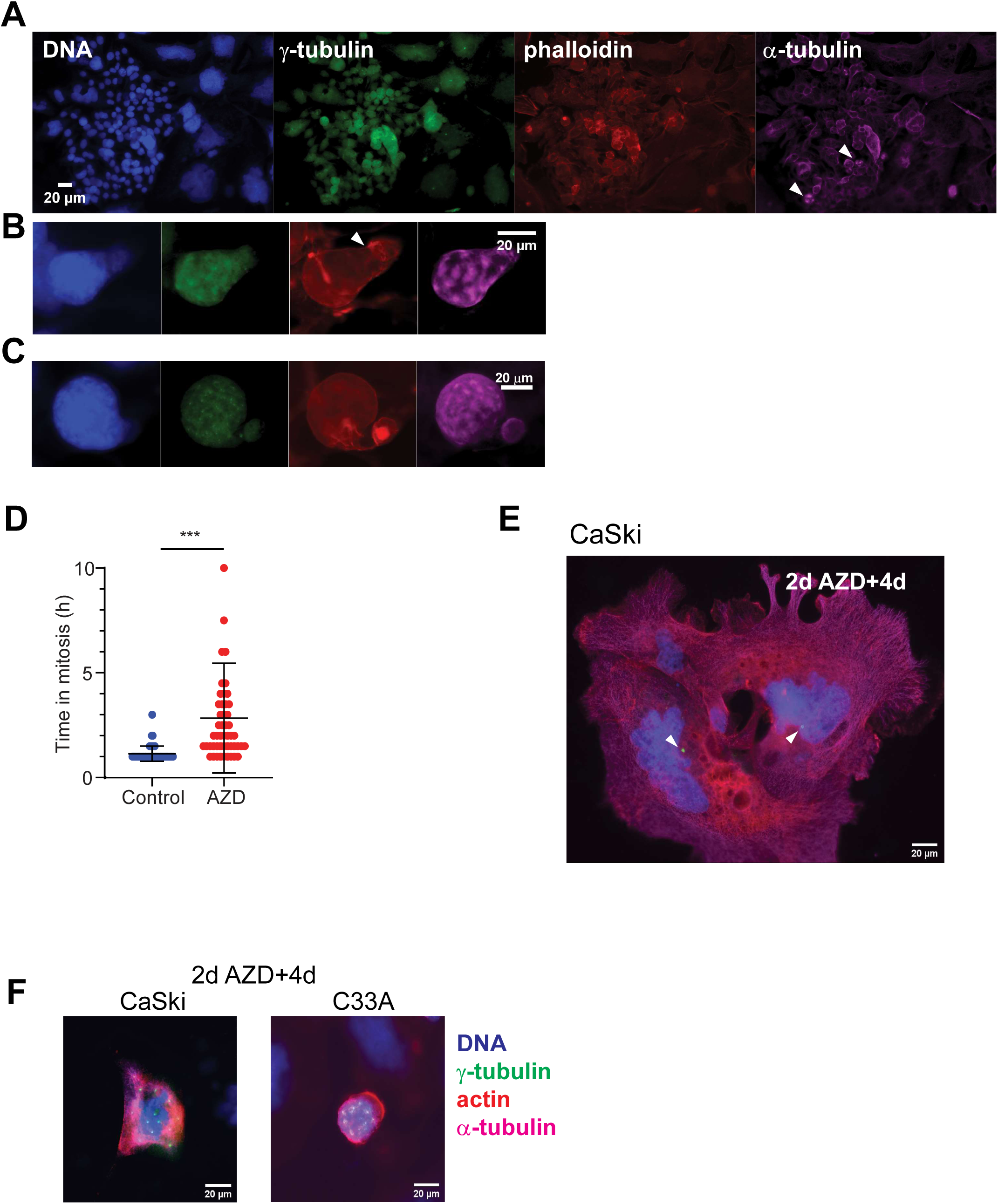
C33A cells treated with 200 nM AZD2811 for 6 days, fixed and stained as in Figure 4. **A:** Small colony of smaller cells with tripolar mitosis (arrowheads). **B:** Large hyper-polyploid mitotic cell with appearance of actin contractile ring (arrowhead). **C:** Large hyper-polyploid mitotic cell with an anucleate bud. **D:** Time of C33A cells treated for 2 days with 200 nM AZD2811 then washed off and followed by time lapse microscopy from 6 – 10 day after drug treatment in mitosis of cells**. E,F**: CaSki and C33A cells were treated with 200 nM AZD2811 for 2 days, then the drug washed out with fresh media and cells were allowed to grow a further 4 days (2d AZD+4d) then fixed and immunostained for the indicated markers.

The time lapse and colony formation experiments were performed on cells that had been incubated with drug for only 2-3 days then the drug was washed out and replaced with fresh media for up to 10 days. The lack of AURKBi resulted in large polyploid cells delaying in mitosis for extended periods (>3 times the average normal mitosis Figure 5D). Immunofluorescence microscopy with the same markers of mitotic spindles as above revealed the same excess of centrosomes clustered in interphase cells and forming multiple spindle poles in mitotic cells as with continuous treatment (Figure 5E,F).

### Tetraploid cells retain long term proliferative potential

A prediction from our experimental data is that tetraploid cells have an increased likelihood of producing proliferative clones whereas once a cell has exceeded 4-8n DNA content and corresponding increased centrosome number, it is improbable that proliferative clones will arise. To test this, RB and p53 defective cells were treated with 200 nM AZD2811 for either 1 (1 failed cytokinesis) or 3 days (>2 failed cytokinesis) then the drug removed and colony formation followed. Even 1 day treatment significantly reduced colony formation in all cell lines, but all were capable of forming small colonies, whereas none were found in the 3 day wash off plates (Figure 6A,B). Analysis of the nuclear DNA content of the individual colonies revealed that the colonies in the 1 day treated cells were on average tetraploid, although the ploidy of different colonies varied considerably (Figure 6C). When the cells were allowed to continue proliferating, it was found that RB^−/−^p53^−/−^ HCT116 cells retained a significant proportion of tetraploid cells that continued to proliferate whereas C33A were predominantly their original ploidy (EdU and Ki67 positive; Figure 6D, Figure S4). This suggest that loss of ploidy is a cell line dependent event but that tetraploidy is well tolerated by cancer cells with loss of RB and p53 function.

**Figure 6.**
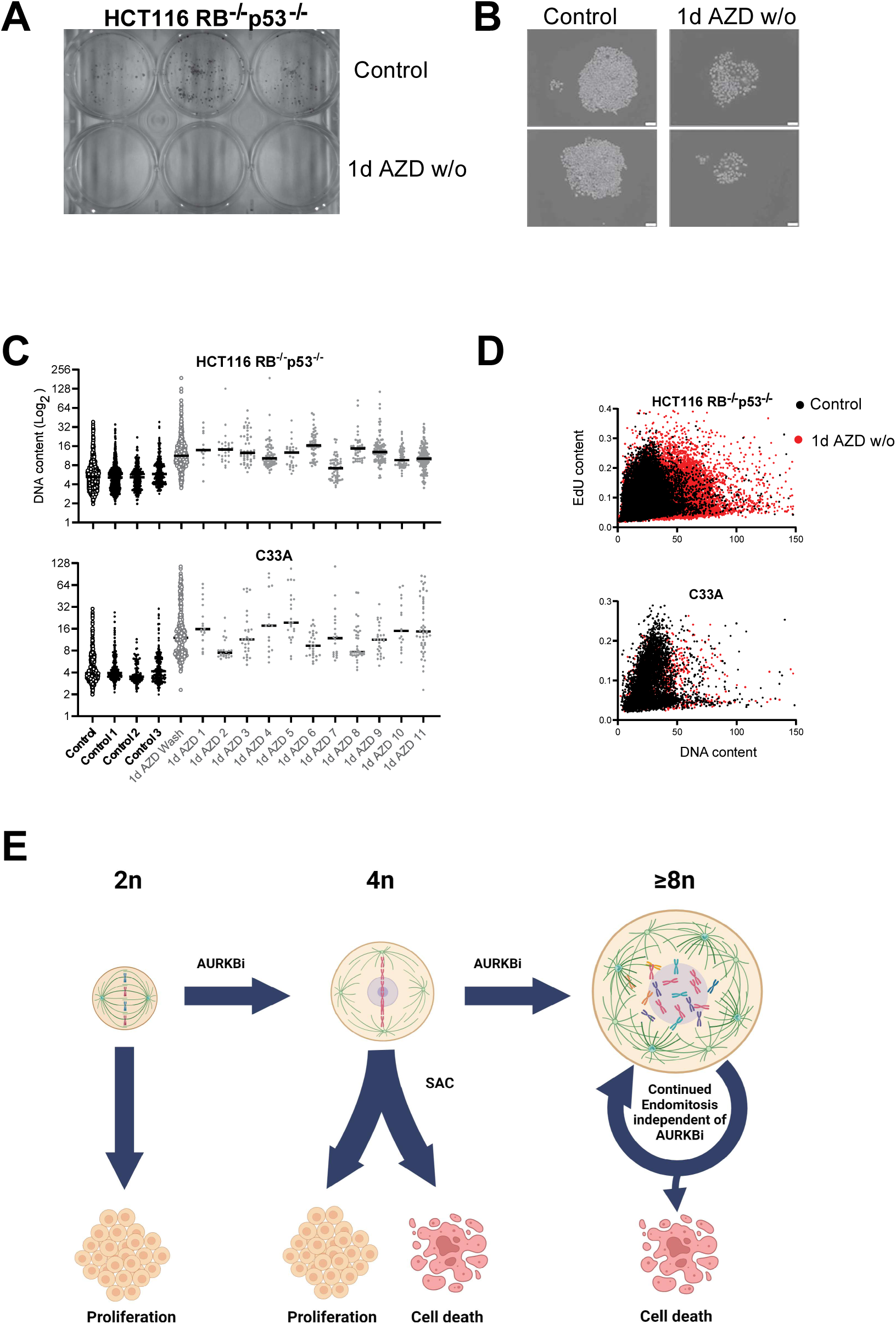
**A**. Colony formation assay of HCT116 RB^−/−^p53^−/−^ control and 1 day AZD2811 treated then washed off cells. **B**. Colonies from control and 1 day AZD-treated cells from A. **C**. DNA content of individual colonies of HCT116 RB^−/−^p53^−/−^ and C33A cells from parallel experiment to A. **D**. Cells grown from the combined colonies in a similar experiment to A, were labelled with EdU, fixed and stained for EdU incorporation and DNA content.

## 3. Discussion

Although loss of RB has been reported to sensitise cells to AURKB inhibitors [24, 29, 32], analysis of the large DepMap data set indicates the majority of RB defective cancer cell lines are relatively unaffected. Inhibition of AURKB can reduce death in cells that have been driven into mitosis with DNA damage by overcoming the spindle assembly checkpoint and driving cells out of mitosis [38], supporting the observation that AURKBi are relatively non-toxic agents, at least in short-term viability assays. Over-expression of the pro-apoptotic BH3-only protein BID has now been identified as a major determinant of increased sensitivity to killing by AURKBi [39]. Loss of RB function will bypass senescence resulting in the large hyper-polyploid cells that we show have a reduced viability over time. It is likely that over-expression of BID further sensitizes the RB-defective large hyper-polyploid resulting from AURKBi treatment to death. However, in the majority of tumour cell lines AURKBi efficiently promotes polyploidy, although the longer-term outcomes of the polyploidy induced differed depending on their RB and p53 status. The loss of RB and p53 function by HPV E6/E7 expression or direct mutation/deletion of RB and p53 blocked ability of AURKBi to promote cell cycle arrest, resulting in endomitotic giant polyploid cells (EPGCs). The data presented here demonstrate that these EPGCs have little DNA damage, despite having extremely altered nuclear morphology with many fractured and micro-nuclei. They continue to undergo DNA replication and mitosis but are incapable of undergoing a productive cytokinesis after at least 2 failed cytokinesis, and incapable of forming viable colonies with long term proliferative potential either *in vitro* or *in vivo*. This distinguishes them from the previously identified polyploid giant cancer cells (PGCCs).

Chemotherapeutic drugs such as doxorubicin that are reported to induce polyploidy *in vitro* cause high levels of DNA damage {Illidge, 2000 #107;Salmina, 2019 #5365;Mosieniak, 2015 #94;Song, 2021 #5587}. The primary response to these drugs is cell cycle arrest followed by death, although a minor proportion of cells can become polyploid [16, 40, 41]. The two outcomes of polyploidy are either senescence or a depolyploidization event which leads to cells that proliferate normally {Mosieniak, 2015 #94}. Microtubule targeting drugs such as paclitaxel, docetaxel and vincristine cause a mitotic arrest and can also lead to the formation of polyploid cells due to mitotic slippage [16, 40, 42]. Long term culture of these polyploid cells produce a mainly diploid population, also suggesting depolyploidization. In the majority of the studies investigating PGCCs, the treatments used to produce them have been either relatively short-term with a single failed mitosis or cytokinesis producing tetraploid cells, or an extended period of treatment (days) that produced extensive cell killing followed the appearance of PGCCs. The PGCCs produced by the short term treatments have been shown to undergo nuclear fragmentation and nuclear budding {Niu, 2016 #5362;Sundaram, 2004 #113; Mosieniak, 2015 #94}, suggesting that these mechanisms may be responsible for the progeny with reduced ploidy. The process of depolyploidization may involve segregation of the replicated genome via multipolar mitosis resulting in nuclear budding or segregation into multiple potentially non-identical daughter cells {Erenpreisa, 2005 #115;Erenpreisa, 2000 #116;Salmina, 2019 #5365}. However, detailed cell fate studies have demonstrated that the most common outcome of multipolar division from either tri– or tetrapolar mitosis is cell cycle arrets or death [37, 43]. We have observed budding and unequal cytokinesis in giant cells 6 days after AURKBi treatment, although *in vitro* colony formation and *in vivo* tumour formation experiments have failed to detect proliferative clones. The mathematical modelling of probability of viable daughter cell from >8 centrosome spindle polar mitosis is exceedingly small [37]matching our experimental observations. An alternative explanation for the viable proliferative cells from longer term treatment is they may represent diploid, slow cycling, drug-tolerant persister cells [44]. These can be reversibly arrested to re-start normal proliferation after drug removal and have acquired enhanced chemo-resistance. These resistant persister rather than depolyploidised PGCCs may be responsible for the claimed attributes of the diploid cells derived from these treatments.

We have demonstrated using 1 day treatment with AZD2811 to produced tetraploid cells that these can form viable colonies and have long term proliferative potential as diploid and tetraploid colonies. This supports the detailed fate studies of enforced tetraploid cells demonstrating reduced long term proliferative potential but stable tetraploid progeny is a potential outcome [37, 45], and the genomics data demonstrating that tetraploidy can be tolerated by tumour cells [9], and In contrast to tetraploid cells, cells that undergo >2 failed cytokinesis and accumulate >8n DNA content and a corresponding increase in centrosome number have lost colony forming ability and long term proliferative potential. Indeed, in many of these experiments, exposure to AURKBi for the initial 2-3 days was sufficient to drive continued endomitotic cycles of failed cytokinesis. It also appears that once cells have failed cytokinesis twice, they will continue to undergo further endomitotic cycles with failed cytokinesis in the absence of AURKBi. The increased centrosomes numbers and their ability to form individual mitotic spindles at each mitosis, rather than cluster to form pseudo-bipolar spindles that have been shown to enable relatively normal mitosis although commonly leading to cell death in subsequent mitosis [36, 43, 45]. The loss requirement for continued AURKBi treatment after 2-3 days is likely to be due to an inability of the mitotic, multipolar (>8 poles), hyper-polyploid (>8n DNA) cells to satisfy the spindle assembly checkpoint and undergo mitotic slippage without cytokinesis. Other studies have indicated the daughters of excess centrosome pseudo-bipolar spindle division are more treatment resistant and form more aggressive tumours, whereas too many centrosomes and spindle poles drive intolerable levels of genomic instability and cell death [46, 47].

AURKB inhibitors have been used in many clinical trials, either as single agents or in combinations [48]. Limited clinical efficacy has been observed in AML [49], but importantly there is no evidence that AURKBi promote more aggressive disease despite evidence the drugs are promoting polyploidy *in vivo* [50]. This matches preclinical studies that have shown in RB and p53 defective tumours that stable disease is the common outcome and the tumours display extensive polyploidy [24].

In summary, AURKBi treatment of RB and p53 defective tumour cells efficiently promotes polyploidy and extended treatment beyond 2 failed cytokinesis is sufficient to drive hyper-polyploidy. These cells, EPGCs, although viable and undergoing continuous endomitotic cycle are incapable of long term proliferative potential either *in vitro* or *in vivo*. It may be possible to use this approach to selectively target RB and p53 defective tumours which represent ∼10% of all tumours, but are >80% of small cell lung cancers and almost all virally driven tumour, driving cells into an EPGC state that may have unique vulnerabilities that can be exploited.

## Materials and Methods

### Cell lines

The parental HCT116 (colorectal carcinoma) and CaSki and C33A cell lines were obtained from American Type Culture Collection (Manassas, VA, USA). HCT116 RB^−/−^p53^−/−^ produced from the original HCT116 p53^−/−^ line [51] using CRISPR CAS9 knockout (Vora et al., in preparation). HT1080 (fibrosarcoma) cell line was purchased from CellBank Australia (Westmead, NSW, Australia). The cell lines were cultured as described previously [29].

Human cervical keratinocytes (HCK) stably expressing TERT [52]parental and HPV E6/E7 expressing were provided by Professor Nigel McMillan from Griffith University (Gold Coast, QLD, Australia) and cultured in Keratinocyte Serum Free Medium (GIBCO) supplemented with Bovine Pituitary Extract (50μg/ml) and Epidermal Growth Factor (5ng/ml) as well as 0.035mM of Pen/Strep and CaCl_2_. The cell cultures were maintained at 37°C, in low oxygen (2% O2 and 5% CO2).

### High Content Imaging

Cells were grown on glass coverslips or in black wall clear bottom 96 well plates (Costar) then treated with AURKBi or DMSO control for the indicated time. Cells were labelled with 10 μM EdU for 2 h and then fixed with 4% PFA, permeabilised with 0.1% Triton X-100 and blocked with 3% BSA. Cells were stained with Cy-5 azide using click reaction for detecting cells that had EdU incorporated into their DNA. Cells were then immunostained using an anti-human Ki-67 (Dako # M724001) antibody and nuclear DNA was stained using 4,6-diamidino-2-phenylindole (DAPI) stain. Plates were imaged using the IN Cell 6500HS imaging system (GE Healthcare), the images were analysed using Cell Profiler version 4.2.1 (Broad institute of MIT and Harvard) and data processed using either GraphPad Prism 8 or R Studio as described previously [53].

### Immunofluorescence

Cells grown on glass coverslips within 6 well plates were fixed, permeabilised and blocked as described. Cells were immunostained with antibodies against either γH2AX (Cell Signalling #2595) or γ-tubulin (Santa Cruz Biotechnology #sc-17787), α-tubulin (Abcam #ab18251) and incubated with phalloidin and DAPI to stain the actin filaments and DNA respectively. Cells were imaged using either the Olympus BX63 upright fluorescent microscope or the Olympus FV3000 confocal microscope.

### Timelapse microscopy

Cells were seeded in a 12-well plate in triplicates and treated with DMSO (control) and AURKi. Live cell imaging was performed using the Zeiss Axio Observer 7 imaging system in 5% CO2 and 37°C and imaged every 20 minutes for 48h-96h. Movies were analysed manually by recording time spent in interphase, mitosis and observing successful divisions, failed cytokinesis or death for each analysed cell.

### Flow cytometry

Cells were harvested and fixed as described previously [29]. The FACS analysis was conducted using the Beckman Coulter CytoFLEX-S. DNA content (PI signal) was analysed ungated, using the PE filter. Data was analysed on Flowjo (version 7.6.4, Becton, Dickinson & Company).

### Colony formation assay

Cells were seeded sparsely into a 6-well plate and treated with either DMSO or AURKBi for either 24h or 72h. After treatment, the cells were washed with PBS and replaced with fresh media and incubated until colonies were observed. Colonies were fixed with 4% PFA and stained with 0.05% Crystal violet. Colonies from control and 1 day treated cells were stained with Hoechst 33372 to record nuclear DNA content prior to crystal violet staining.

### Mouse tumour assay

Animal experiments were performed with approved ethics from the The University of Queensland Animal Ethics Committee (2022/AE000024). 6-week old Nude Mice (ARC) were injected with 3 x 10^6^ control or 5d AURKBi treated cells in Matrigel. Tumour free survival was measured and the mice were culled when tumours were palpable.

### Analysis of DepMap data

Analysis of both the GDSC2 [35]and CTD^2 [34] small molecule viability datasets in DepMap was performed using the DepMap portal Data Explorer (http://www.depmap.org). Alisertib (MLN8237) was present in both datasets (725 cell lines in CTD^2; 404 cell lines in GDSC2), Barasertib (AZD2811) was present in the CTD^2 (725 cell lines). The area under the inhibition curves (AUC) was used for each dataset, with lower AUC indicating sensitivity to the drug. Expression, mutation, copy number and antibody staining intensity for all the cell lines was derived from the Cancer Cell Line Encyclopedia (CCLE; [54]). The sensitivity to Barasertib and Alisertib were highly correlated in the CTD^2 datasets (Spearman r = 0.67), as was the sensitivity to Alisertib in the cell lines overlapping between the CTD^2 and GSCD1 set (Spearman r = 0.46).

## Author Contributions

Conceptualization, BG, NAJM; methodology, SV, AA, RPK, MF, YH; formal analysis, BG; investigation, SV, AA, RPK, DN, BG; resources, YH, JH, NAJM, JU, BG; writing—original draft preparation, SV, AA, BG; writing—review and editing, SV, AA, RPK, DN, MJKJ, YH, JH, NAJM, JU, BG; visualization, BG.; supervision, BG; project administration, BG; funding acquisition, NAJM, BG. All authors have read and agreed to the published version of the manuscript.

## Acknowledgments

The authors thank Prof Nikolas Haass for useful discussions on interpretation and Dr Martina Proctor for assistance with cell lines.

## Figure Legends

**Supplementary Figure S1.**
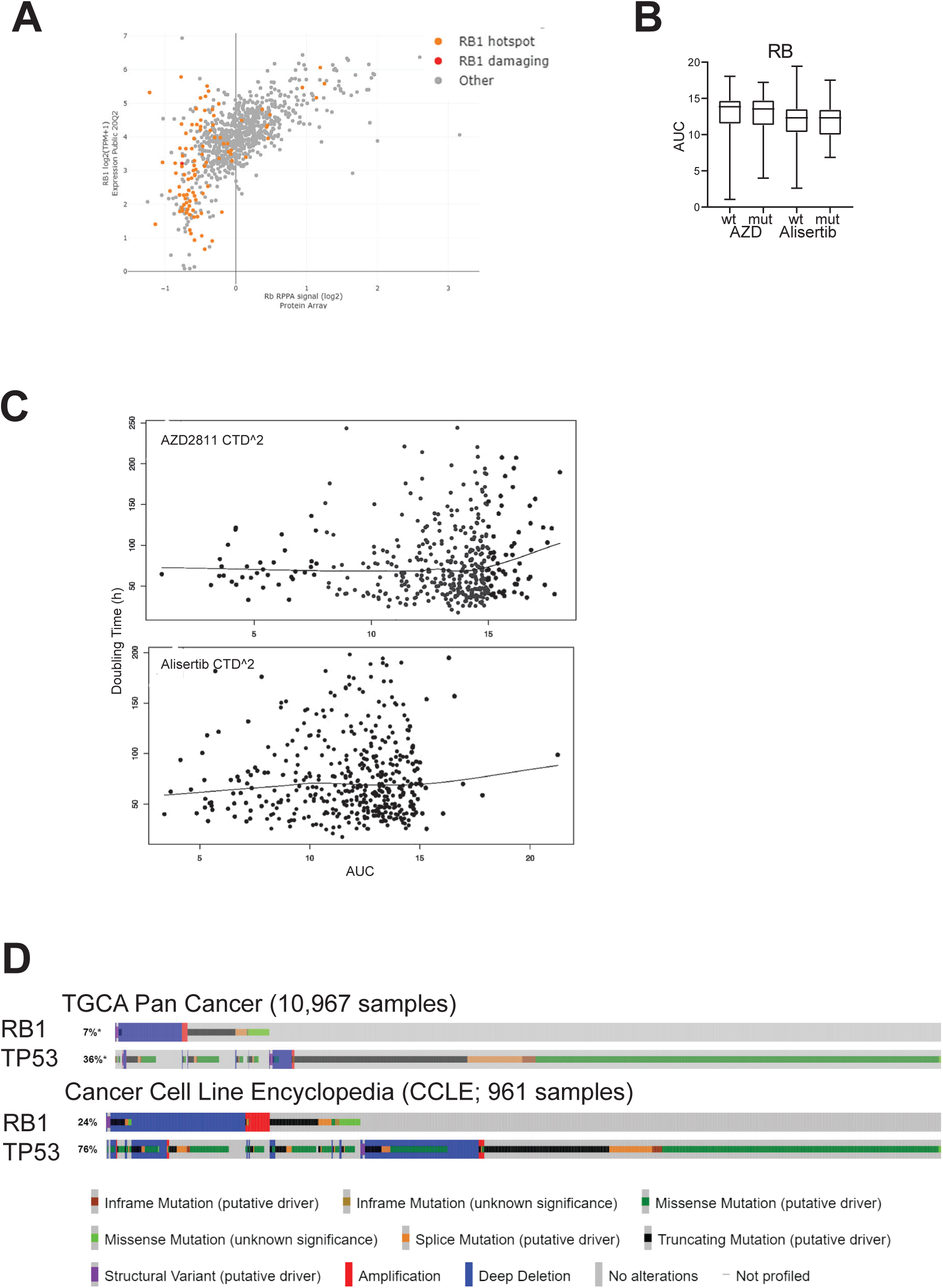
**A**: Expression data for *RB1* showing RNAseq and reverse phase protein array analysis of the cell lines analysed in the CTD^2 screen. The orange and red dots are mutant *RB1*. **B**. The DepMap sensitivity data for AZD2811 (AZD) and Alisertib (Alis) from the CTD^2 drug screen was analysed on the basis of presence or absence of RB mutation. The AUC values (area under the dose response curve) was used to determine sensitivity, the higher values indicate less sensitivity. The box and whiskers plot show the AUC for each drug for all 729 cell lines analysed. **C**. Doubling time v the sensitivity to AURKBi for cell lines in the CTD^2 screen. The line shows the mean of the doubling times. **D.** Mutation and depletion analysis of RB1 and TP53 from TGCA and CCLE databases analysis using cBioPortal.

**Supplementary Figure S2.**
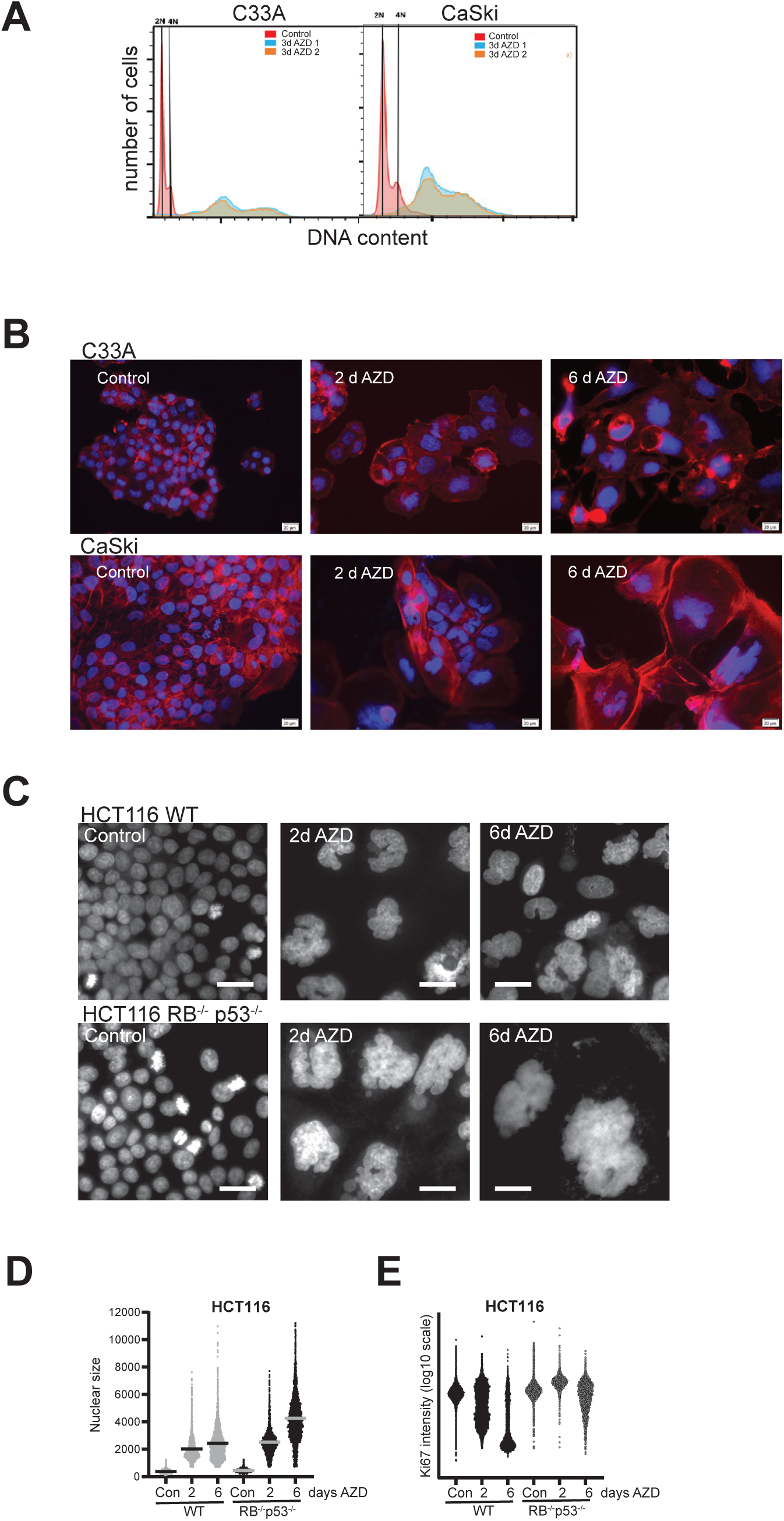
**A**: C33A and CaSki cells were treated with 200 nM AZD2811 for 3 days, fixed and stained for DNA, then analysed by flow cytometry. Control cells are shown as the red histogram, and replicate AZD2811 treated cultures are shown as blue and orange histograms. **B:** C33A and CaSki cells were treated with 200 nM AZD2811 for the indicated time, fixed and stained for DNA and actin, then imaged. The bar indicated 20 μm. **C:** HCT116 RB and p53 wild type (WT) and RB and p53 knockout cells were treated with 200 nM AZD2811 for the indicated time, fixed and stained for DNA then imaged. The bar indicated 50 μm. **D,E:** The indicated genotype of HCT116 cells were treated as in A, fixed and stained for DNA and Ki67 then analysed using high content imaging. The nuclear size and Ki67 intensity are shown. In each case >4, 000 cells were analysed.

**Supplementary Figure S3.**
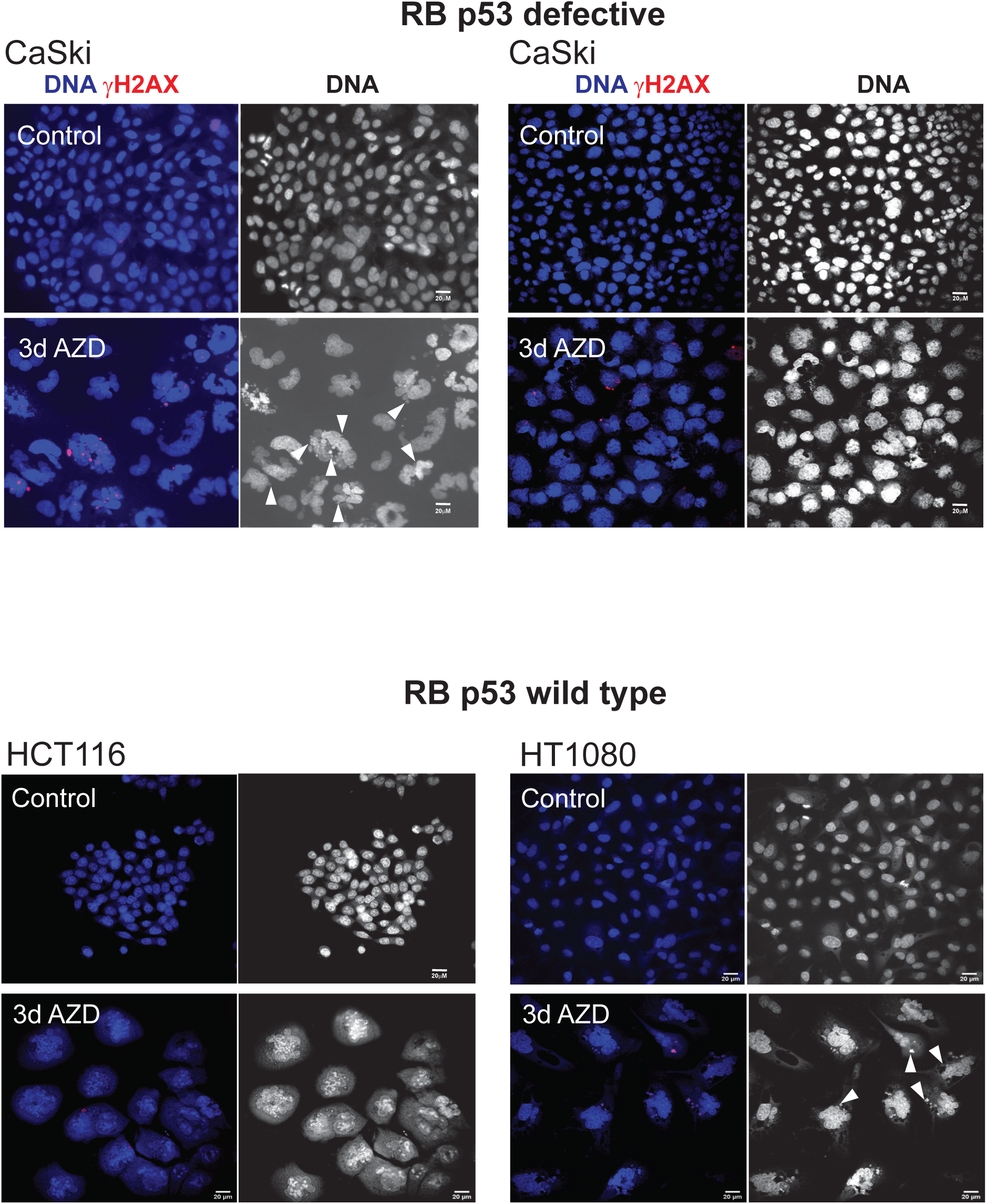
RB and p53 defective CaSki C33A cells RB and p53 wild type HCT116 and HT1080 cells, control and treated with 200 nM AZD2811 for 3 days, fixed and stained for DNA and γH2AX. γH2AX stained micronuclei are indicated (arrowheads).

**Supplementary Figure S4.**
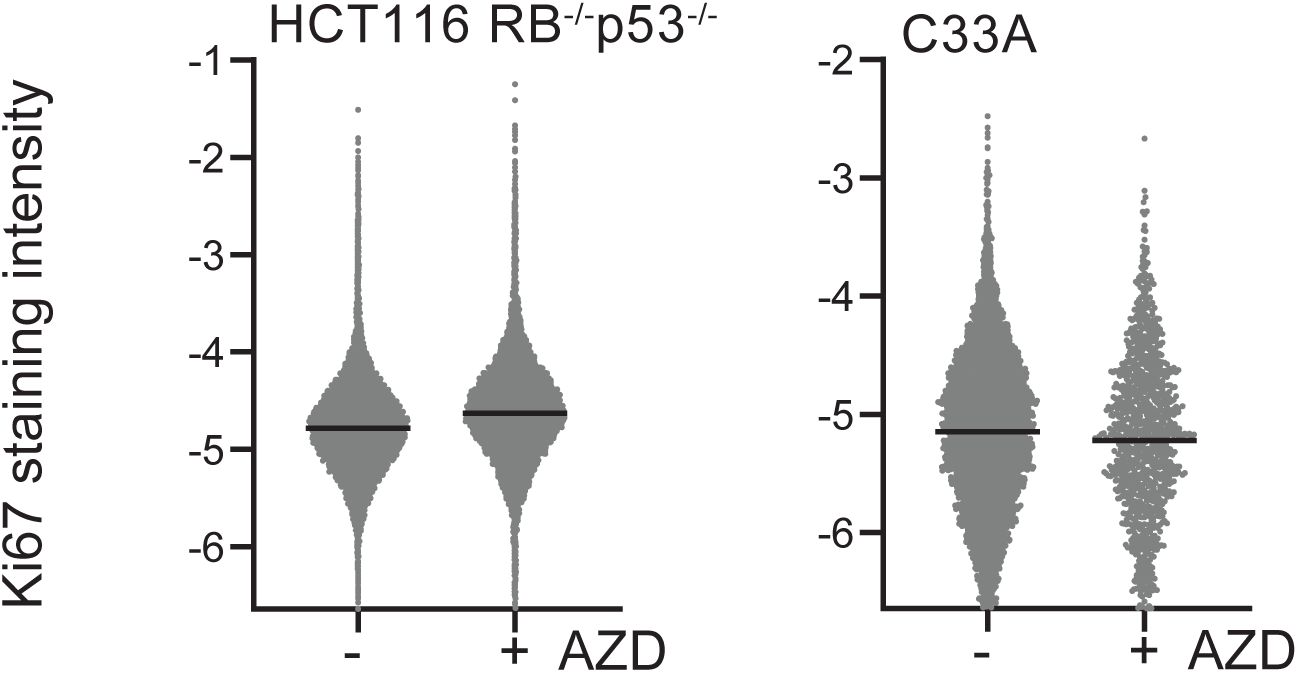
Ki67 staining intensity of cell lines derived from either HCT116 RB-/−p53-/− or C33A line, either control or treated for 1 day with 200 nM AZD2811 then washed out and allowed to grow in fresh media as in Figure 6C.

